# Elucidating the heterogeneity of immunotherapy response and immune-related toxicities by longitudinal ctDNA and immune cell compartment tracking in lung cancer

**DOI:** 10.1101/2023.06.23.546338

**Authors:** Joseph C. Murray, Lavanya Sivapalan, Karlijn Hummelink, Archana Balan, James R. White, Noushin Niknafs, Lamia Rhymee, Gavin Pereira, Nisha Rao, Jillian Phallen, Alessandro Leal, David L. Bartlett, Kristen A. Marrone, Jarushka Naidoo, Benjamin Levy, Samuel Rosner, Christine L. Hann, Susan C. Scott, Josephine Feliciano, Vincent K. Lam, David S. Ettinger, Qing Kay Li, Peter B. Illei, Kim Monkhorst, Ali H. Zaidi, Robert B. Scharpf, Julie R. Brahmer, Victor E. Velculescu, Patrick M. Forde, Valsamo Anagnostou

## Abstract

**Purpose:** Although immunotherapy is the mainstay of therapy for advanced non-small cell lung cancer (NSCLC), robust biomarkers of clinical response are lacking. The heterogeneity of clinical responses together with the limited value of radiographic response assessments to timely and accurately predict therapeutic effect -especially in the setting of stable disease-call for the development of molecularly-informed real-time minimally invasive predictive biomarkers. In addition to capturing tumor regression, liquid biopsies may be informative in evaluating immune-related adverse events (irAEs).

**Experimental design:** We investigated longitudinal changes in circulating tumor DNA (ctDNA) in patients with metastatic NSCLC who received immunotherapy-based regimens. Using ctDNA targeted error-correction sequencing together with matched sequencing of white blood cells and tumor tissue, we tracked serial changes in cell-free tumor load (cfTL) and determined molecular response for each patient. Peripheral T-cell repertoire dynamics were serially assessed and evaluated together with plasma protein expression profiles.

**Results:** Molecular response, defined as complete clearance of cfTL, was significantly associated with progression-free (log-rank p=0.0003) and overall survival (log-rank p=0.01) and was particularly informative in capturing differential survival outcomes among patients with radiographically stable disease. For patients who developed irAEs, peripheral blood T-cell repertoire reshaping, assessed by significant TCR clonotypic expansions and regressions were noted on-treatment.

**Conclusions:** Molecular responses assist with interpretation of heterogeneous clinical responses especially for patients with stable disease. Our complementary assessment of the tumor and immune compartments by liquid biopsies provides an approach for monitoring of clinical benefit and immune-related toxicities for patients with NSCLC receiving immunotherapy.

**Statement of translational relevance:** Longitudinal dynamic changes in cell-free tumor load and reshaping of the peripheral T-cell repertoire capture clinical outcomes and immune-related toxicities during immunotherapy for patients with non-small cell lung cancer.

## Introduction

Despite the routine use of immunotherapy for patients with metastatic non-small cell lung cancer (NSCLC), intrinsic and acquired resistance to immunotherapy regimens are common. Although PD-L1 expression guides selection of first-line therapy, it does not always predict clinical response (1). A multitude of tumor and immune biomarkers – from tumor mutational burden (TMB) to immune-related gene expression signatures – have been evaluated as predictors of immunotherapy response (2,3), however, with the exception of microsatellite instability most of these putative biomarkers fail to consistently predict outcomes. Optimal patient selection for immunotherapy represents a critical challenge that is intensified by the heterogeneity of clinical responses and inefficiency of imaging to fully and timely capture therapeutic response. As such, for the majority of patients treated with an immunotherapy-containing regimen – representing >70% of patients with NSCLC – clinical equipoise remains between the available options, even with preferences based on histology and PD-L1 expression (4).

Minimally invasive liquid biopsy approaches have been shown to enable early and accurate therapeutic response assessment and long-term response monitoring during immunotherapy. An increasing number of studies have shown that monitoring of early on-therapy ctDNA kinetics, using the mean (5,6) or maximum mutant allele fraction (MAF) (7,8) of tumor-derived alterations can enable real-time tumor burden assessment and have supported the role of ctDNA molecular response as an early endpoint for response to immunotherapy (9–13). Collectively, these studies support the clinical utility of ctDNA response monitoring for immunotherapy-treated NSCLC and importantly highlight areas where ctDNA evaluation can be particularly informative and require further investigation. Given the challenges with imaging to capture the timing and magnitude of therapeutic response in the context of immunotherapy (14), ctDNA dynamics may be particularly informative in identifying patients with radiographically stable disease that have differential outcomes. To date, such associations have only been evaluated in a small number of studies in lung cancer including few patients with radiologic stable disease (11,15). Previous studies have also highlighted the value of longitudinal ctDNA response monitoring throughout the treatment course (8,15–17), which would increase the clinical sensitivity of ctDNA response for predicting therapeutic response.

In addition to identification of patients most likely to attain clinical benefit and those at risk for disease progression, opening a therapeutic window of intervention for the latter and sparring unnecessary interventions for the former, minimally invasive blood-based analyses could be valuable for prediction of emergence of immune related adverse events (irAE) (18–20). Baseline T cell characteristics, including the abundance of activated CD4+ memory T cells and the diversity of the T cell receptor (TCR) repertoire have been linked to the development of severe irAEs (18). Similarly, multiple studies have evaluated immune cell subsets or cytokines/chemokines as predictors of irAEs (15,21–23); however, no prospectively-evaluated biomarkers are used in clinical practice today. Although recent studies have shown a correlation between early clonal expansions of T cell populations and the timing and severity of irAEs, on-treatment biomarkers for therapeutic response assessment and prediction of irAEs to immunotherapy-based regimens remain limited (18).

To study the heterogeneity of immunotherapy response together with the value of peripheral blood immune repertoire analyses in capturing irAEs, we designed a longitudinal study of patients with NSCLC that received IO-based therapies. We assessed ctDNA, peripheral TCR dynamics and plasma protein expression profiles from serial blood specimens to capture timing, depth and quality of tumor and immune responses in the context of immunotherapy.

## Methods

### Cohort characteristics

A total of 30 patients were selected for inclusion in the study using the following criteria: (i) advanced or metastatic NSCLC who received at least one cycle of immunotherapy or chemo-immunotherapy treatment; (ii) at least two plasma samples evaluable for cell-free DNA sequencing; (iii) at least one buffy coat sample for WBC genomic DNA sequencing and/or with either clinical tumor mutational profiling (TMP) or whole exome sequencing (WES) of tumor tissue; and (iv) had clinical follow-up including imaging through to time of death or at least 6 months from time of treatment initiation. Twenty-four patients treated with standard-of-care pembrolizumab (n=13) or chemotherapy with pembrolizumab (n=11) were identified from the Immunobiology Blood and Tissue Collection of Upper Aerodigestive Malignancies biospecimen collection protocol and registry (IRB00100653 approved by the Johns Hopkins University IRB). This included 1 patient from a prior study (Anagnostou et al., Cancer Res, 2019) (15) from whom we collected and analyzed additional samples from 3 on-treatment timepoints (Supplementary Table S1). Six patients were enrolled at the Netherlands Cancer Institute (NKI), in accordance with institutional regulatory approvals and received second-line single agent immunotherapy with nivolumab. Clinical data were abstracted from electronic health records and clinical tissue molecular profiling (TMP) data, including variants, were retrieved from available clinical next-generation sequencing outputs. PD-L1 expression, assessed by immunohistochemistry, was abstracted from medical records and evaluated as follows: high expression of PD-L1 on tumors was defined as ≥50% tumor cell staining, and low or negative expression was defined as <50% tumor cell staining. Primary toxicities attributed as immune related adverse events (irAE) were assessed from medical records according to the CTCAE version 5.0 grading system (24) (Supplementary Table S1). Radiographic response assessments were retrieved from the medical records. Progression-free and overall survival (PFS and OS) were assessed as well as durable or non-durable clinical benefit (DCB or NDB). DCB was defined as lack of progression or death at 6 months; NDB was defined as progression or death within 6 months from immunotherapy initiation. Studies were conducted in accordance with the Declaration of Helsinki, approved by the Institutional Review Board (IRB), and patients provided written informed consent for samples and clinical data for research purposes.

### Plasma cell-free, WBC and tumor next generation sequencing (NGS)

Plasma and whole blood samples were collected at times when standard-of-care clinical venipuncture was indicated and processed for targeted error-correction sequencing (TEC-Seq) as previously described (15). Cell free DNA was isolated from plasma using the Qiagen Circulating Nucleic Acids Kit (Qiagen GmbH). For 29 patients, matched white blood cell (WBC) DNA was extracted from buffy-coat obtained at baseline and/or after treatment initiation, using the Qiagen DNA Blood Mini kit (Qiagen GmbH) and sheared to a target fragment size of 200bp. TEC-Seq DNA libraries were prepared from 6.5-125ng cfDNA and 75ng WBC DNA, followed by targeted capture using a custom set of hybridization probes (PlasmaSELECT-63 or -64 panels, Personal Genome Diagnostics [PGDx], Baltimore, MD; Supplementary Table S2) and sequenced using 100 bp paired-end runs on Illumina HiSeq 2000/2500 instruments. TEC-Seq characteristics are described in Supplementary Table S3.

Matched tumor/normal whole exome sequencing (WES) was performed for 17 patients and WES data was used to identify tumor-specific mutations in plasma. Briefly, tumor DNA was extracted following pathological review and macro-dissection of formalin-fixed paraffin-embedded (FFPE) tumor tissue using the Qiagen DNA FFPE kit (Qiagen GmbH). Tumor and matched WBC DNA were sheared to a target fragment size of 200bp and processed for WES as described previously (8). Hybrid capture of exonic regions was performed using Agilent SureSelect in-solution capture reagents and SureSelect XT Human All Exon V4 probes (Agilent). Captured libraries were sequenced using 100 bp paired-end runs on Illumina HiSeq 2500. WES technical metrics are described in Supplementary Table S4.

### Variant calls, annotation, and origin classification

TEC-Seq data was analyzed as previously described using a genomics annotation pipeline including: primary processing by Illumina CASAVA software (v1.8), alignment to the human reference genome (hg19/GRCh37) by Novoalign, and variant calling by VariantDx (25). Supermutant counts were defined as the total number of barcoded read families in which ≥95% of members contained the same variant. Variants in plasma cfDNA were defined as detectable at a supermutant count of at least 3, whereas variants in WBC specimens –that were also detected in plasma cfDNA-were defined as detectable at a supermutant count of at least 1. The variant allele fraction (VAF) for a given plasma variant was determined from the percentage of distinct mutant versus total cfDNA variant reads at a given locus.

The origin of plasma cfDNA variants was classified using all available sequencing data, including matched tumor (clinical TMP or WES) and all WBC DNA sequencing, as tumor-derived, germline or clonal hematopoiesis (CH)-derived, based on a tiered approach. Patients with matched tumor next-generation sequencing (NGS) and detectable tumor-specific mutations in plasma (n=21) were evaluated using a tumor-informed approach for the identification of ctDNA variants. Given the limitations of single-region tissue NGS for capturing the molecular heterogeneity of individual tumors, patients with matched tissue sequencing but no detectable tumor-specific mutations in plasma (n=3) were further evaluated using a WBC-informed approach (8) to identify any candidate ctDNA variants missed by tissue NGS. Patients with no detectable tumor-derived mutations in plasma according to both approaches were subsequently classified as undetectable. Similarly, for 6 patients without matched tumor tissue NGS available (n=6), we used the WBC-informed approach to characterize variant origin in plasma and cases with no detectable tumor-derived mutations were classified as undetectable (n=1). For patients with multiple WBC DNA samples sequenced (n=11), the intersection of all alterations detected across sequenced WBC samples was used.

Plasma variants were cross-referenced against the Catalogue of Somatic Mutations in Cancer (COSMIC) v95 database for the annotation of hotspot alterations using OpenCRAVAT (26). A conservative COSMIC frequency threshold of 25 hits was used to define a lung cancer hotspot with high confidence (8,27). Of the variants that were not classified as a hotspot, those with a variant allele frequency ≥25% in all plasma and available WBC samples from a patient were classified as germline. Plasma variants detected in genes commonly altered in clonal hematopoiesis-CH, namely *DNMT3A, TET2, ASXL1, PPM1D, TP53, JAK2, RUNX1, SF3B1, SRSF2, IDH1, IDH2, U2AF1, CBL, ATM and CHEK2* (CH blacklist; Supplementary Table S5) and a supermutant count ≥1 in matched WBC TEC-Seq were considered CH-derived. Plasma variants that were not detected in matched WBC sequence data or could not be evaluated as matched WBC sequencing was not performed were classified as follows: variants within a CH blacklisted gene and an alteration frequency ≥5% in COSMIC hematologic malignancies were classified as CH-derived, providing that they were not lung cancer hotspots or reported in matched tumor tissue sequencing. Plasma variants that could not be verified by either matched tumor tissue sequencing or as lung cancer hotspots, and variants that were classified as a lung cancer hotpot within a CH-blacklisted gene, were assigned as variants of unknown origin (VUO). For WBC-informed classifications, all non-germline variants were further compared against matched WBC DNA sequence data to determine cellular origin. Plasma variants within a CH-blacklisted gene and a supermutant count ≥1 in matched WBC were considered CH-derived. Given the small possibility of buffy-coat contamination by circulating tumor cells, resulting in the misclassification of tumor-derived alterations as CH-derived, hotspot variants that were detected in matched WBC within a non-CH blacklisted gene were classified as tumor-derived. Plasma variants within a CH-blacklisted gene and an alteration frequency ≥5% in COSMIC hematologic malignancies were classified as CH-derived providing that they were not lung cancer hotspots. A summary of the branched logic architecture used for variant origin classification is shown in Supplementary Figure S1.

### Cell-free tumor load tracking and molecular response evaluation

The mutant allele fraction (MAF) of the most abundant tumor-derived mutation at each serial plasma sample (referred to as the maximum MAF) was used as a measure of circulating cell-free tumor load (cfTL) and a reduction of cfTL to 0% indicated cfTL clearance (8). Patients who had complete cfTL clearance from baseline to the final timepoint analyzed were classified in the molecular response category. Patients with cfTL clearance followed by a subsequent increase were classified in the molecular response followed by recrudescence category. Patients with cfTL persistence across all timepoints analyzed were assigned a classification of molecular progression.

### TCR sequencing

TCR Vβ next-generation sequencing was performed for 30 patients (comprised of 23 patients included in cfTL analyses and 7 additional patients from Anagnostou et al., Cancer Res, 2019) from WBC DBA isolated from serial peripheral blood samples prior to treatment initiation (n=23) and during the course of treatment (n=58). Briefly, genomic DNA was isolated and TCR-β CDR3 regions were amplified using the deep ImmunoSeq assay by multiplex PCR specific to TCR Vβ, Dβ, and Jβ regions (Adaptive Biotechnologies). Productive TCR clonotypes were identified and analyzed at the amino acid sequence level. TCR clones were categorized as increasing or decreasing between baseline and each on-treatment timepoint and relative percentages of these clones were compared using Fisher’s exact test with P-value adjustment using the False Discovery Rate (FDR; Benjamini-Hochberg procedure). This was followed by aggregation of clone dynamic measures per subject and statistical comparisons of patient subgroups of interest using the nonparametric Mann-Whitney U test. Unique TCR-β CDR3 sequences were clustered according to anticipated specificity for epitope recognition and HLA-allele associations using the GLIPH2 (Grouping of Lymphocyte Interactions by Paratope Hotspots 2) method (28). GLIPH2 identified global TCR-β clusters based on the similarity of CDR3 sequences as follows: TCRs that shared the same length CDR3 differing by up to seven amino acids with matching global motifs ≥ 4 amino acids in length were grouped into the same cluster. Clusters identified by GLIPH2 were considered for statistical analyses based on the following parameters: detected in ≥ 3 distinct patients, number of unique CDR3 ≥ 2, vb_score (enrichment of V genes within cluster) < 1, length_score (CDR3 length distribution within cluster) < 0.05.

### Plasma proteomics

Expression levels of 92 proteins in the Olink Immuno-Oncology panel (Olink Proteomics AB, Uppsala, Sweden) were measured in plasma samples (n=88) from 28 patients using a proximity extension assay (PEA). Quality control and data normalization were performed based on internal and external controls. Internal controls consisting of an immuno-based non-human antigen control, an extension control comprised of an antibody conjugated to a unique oligonucleotide DNA pair for immediate proximity-dependent hybridization and extension, and a detection control comprised of a synthetic double-stranded DNA amplicon to monitor amplification were spiked into each sample well to assess intra-assay variability (Olink Proteomics AB, Uppsala, Sweden). External sample controls, used to evaluate assay precision, were also included on the assay plate and consisted of negative controls (buffer-only) and inter-plate controls (pool of 92 antibodies conjugated to each oligonucleotide pair). Protein expression values were generated as log2 of Normalized Protein Expression (NPX) as follows: counts of known barcode sequences were first normalized to the extension control, followed by log2 transformation and intensity normalization against the inter-plate control. Mean protein expression values were calculated for each sample, as well as the difference in values between timepoints and compared across different patient groups using the Mann-Whitney U test.

### Statistical analyses

For survival analyses, median point estimates were estimated using the Kaplan-Meier method and survival curves were compared by log-rank testing. Univariate and multivariate Cox proportional hazards regression analyses were used to evaluate the impact of molecular responses on overall and progression-free survival. For categorical comparisons, Fisher’s exact test was utilized. For non-parametric comparisons, the Mann-Whitney U test was utilized. Unless otherwise specified, differences across comparisons were considered significant at p less than 0.05. All analyses were conducted using R Statistical Software (version 4.1.2) as described above (29).

### Data availability

Next-generation sequence data can be retrieved from the European Genome-Phenome Archive (EGA; study accession number EGAS00001007291).

## Results

### Cohort description and plasma variant characterization

We performed deep targeted error-correction sequencing (TEC-seq) of 162 serial plasma cell-free DNA (cfDNA) and 47 white blood cell (WBC) genomic DNA (gDNA) samples from a real-world cohort of 30 patients with advanced or metastatic NSCLC treated with immunotherapy or chemo-immunotherapy regimens (Fig. 1). Plasma cfDNA sequencing was completed from baseline samples prior to treatment in 26 patients (median 0 weeks, range -0.7-0 weeks; Supplementary Table S3). When baseline samples were insufficient or unevaluable (n=4 patients), the next available samples were assessed, prior to the second treatment cycle (median 3 weeks, range 3-3.14 weeks; Supplementary Table S3). Sequence alterations were analyzed in plasma at baseline and longitudinally after treatment initiation.

**Fig. 1.**
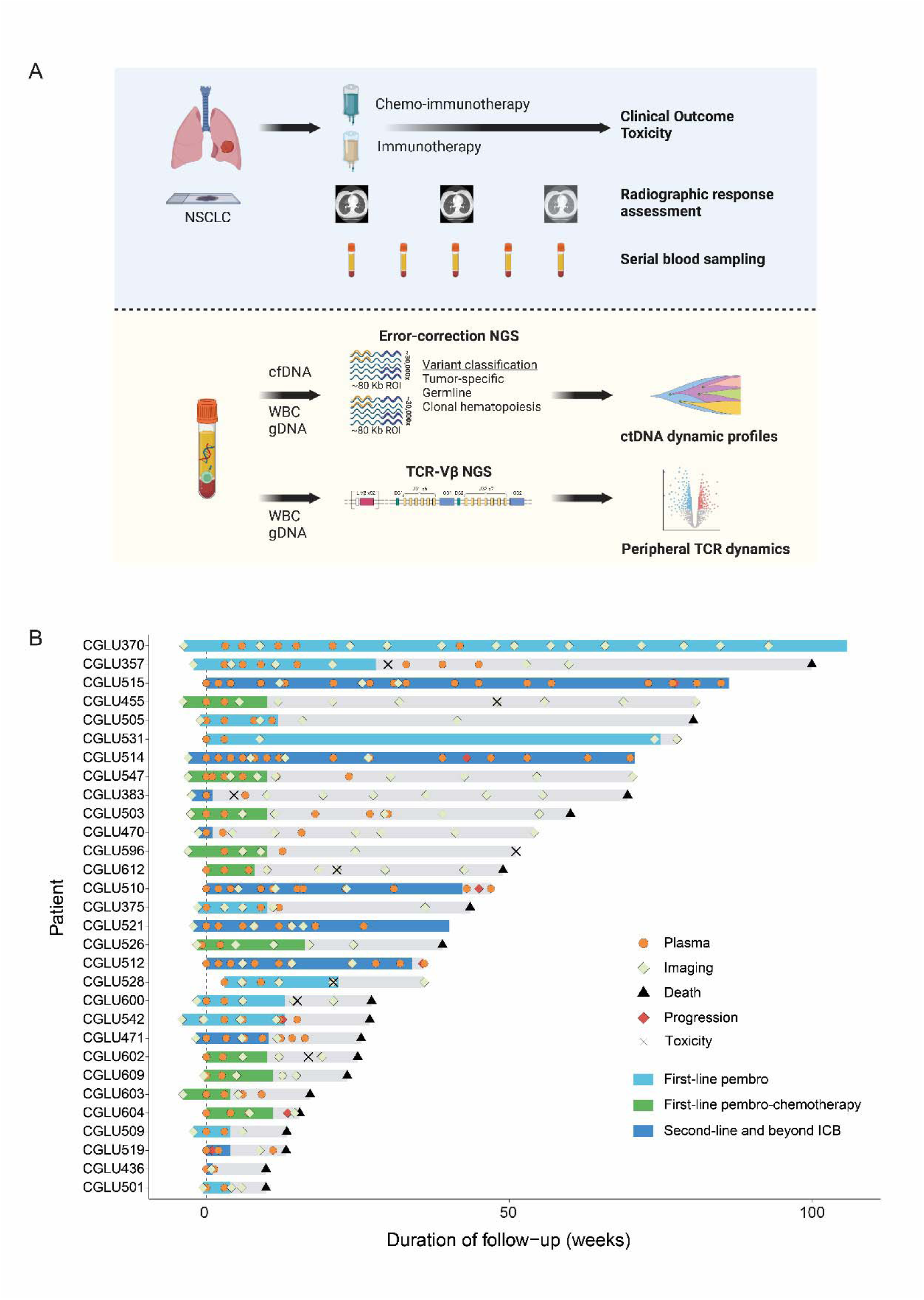
Overview of study methodology and patient cohort. (A) From a biospecimen repository of serial blood collections for patients with advanced non-small cell lung cancer (NSCLC) treated with chemo-immunotherapy or immunotherapy, we assessed clinical outcomes including toxicity and radiographic response. High-depth targeted error-correction sequencing (TEC-Seq) was performed on plasma cell-free DNA (cfDNA) and available matched white blood cell (WBC) genomic DNA samples from 30 patients. Tumor tissue whole exome sequencing was performed on baseline tumors and buffy-coat samples from 17 patients, and clinical tumor next-generation sequencing (NGS) data was available for 15 patients. The origin of plasma variants was classified using either a tumor- or WBC-informed approach, based on the availability of sequencing data. Longitudinal changes in total cell-free tumor load (cfTL) were tracked for each patient and used for assessment of molecular response to therapy. For a subset of patients, serially collected WBC genomic DNA samples underwent T-cell receptor (TCR) variable-beta (Vβ) chain sequencing for analysis of peripheral TCR dynamics. (B) Swimmer plot showing the course of disease for each patient included in cfTL analyses, alongside treatment and samples analyzed. Radiographic response assessments recorded up to 4 weeks prior to treatment initiation are shown. ICB, immune checkpoint blockade.

A total of 461 variants in 37 genes were detected in plasma cfDNA (Fig. 2A, Supplementary Table S5). The cellular origin of cfDNA variants was determined using all available plasma, WBC DNA and tumor tissue sequencing data according to a tiered approach (Methods); using this classification approach, tumor-derived sequence mutations (43% of all variants with resolved cellular origin) were identified in 26 patients (Fig. 2B, Supplementary Table S5, Supplementary Fig.S2A). This included 2 patients with no detectable tumor tissue-specific variants in plasma, in whom we identified KRAS G12C, G12V and TP53 G245C ctDNA mutations using a WBC-informed approach (Methods). Notably, 25 mutations (6% of all variants with resolved cellular origin) were of germline origin and 205 variants (51% of all variants with resolved cellular origin) were CH-derived (Fig. 2C-D, Supplementary Fig.S2A). CH variants were detected across genes both canonically and non-canonically associated with clonal hematopoiesis, including DNMT3A, TP53, BRCA2 and EGFR (Fig. 2C, Supplementary Table S5) and were removed from downstream analyses of molecular response.

**Fig. 2.**
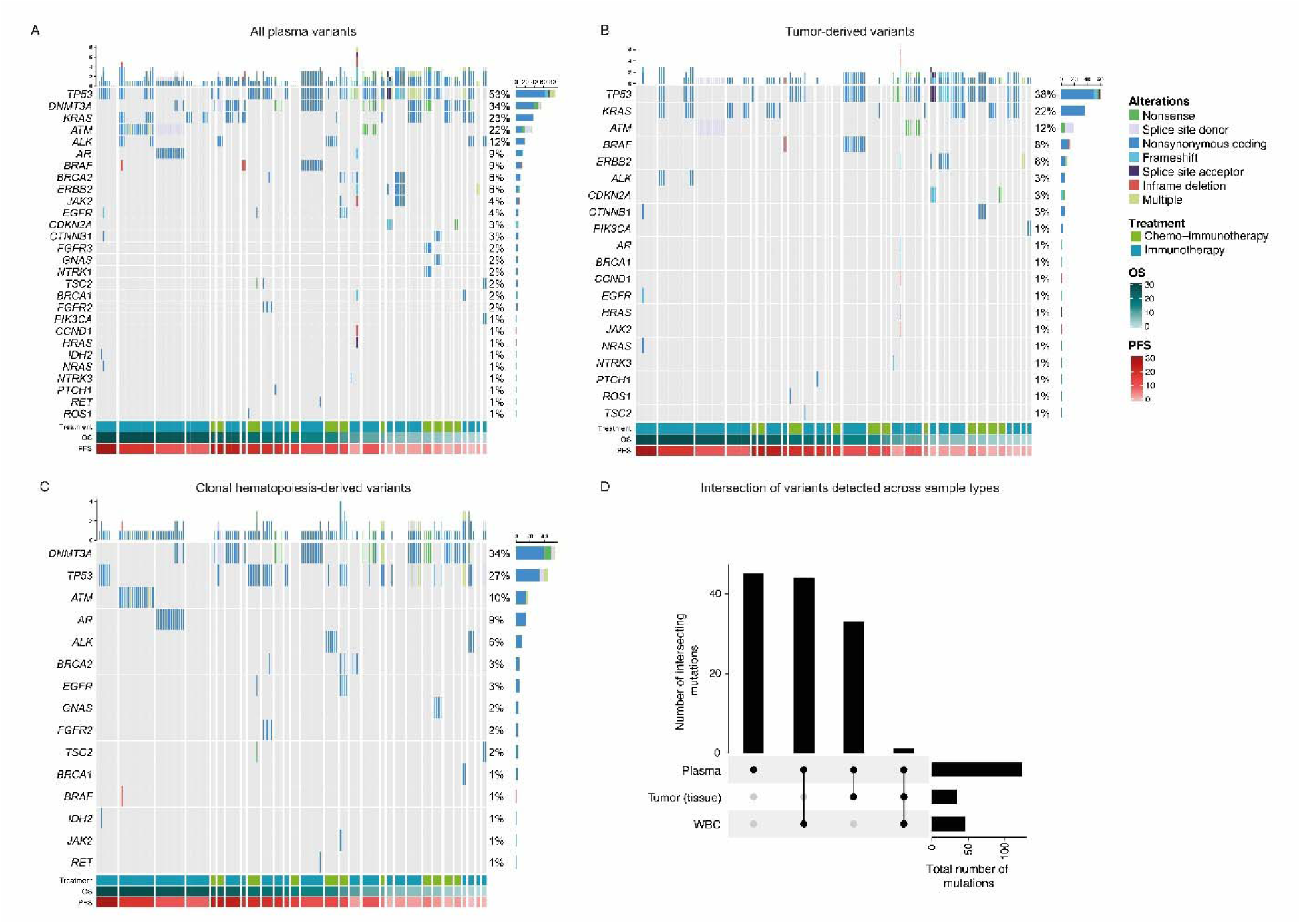
Landscape of variants detected in plasma. (A) Plasma cell-free DNA (cfDNA) variants detected across samples analyzed from all 30 patients in the study cohort, using TEC-Seq. The types of alterations detected across plasma samples are shown alongside alteration frequencies for each gene (right). Per-sample mutation counts are displayed in bar plots on the top. Panels indicating treatment type, overall (OS) and progression-free survival (PFS) are displayed below. (B) Plasma variants identified as being of tumor-derived origin. (C) Clonal hematopoiesis-derived white blood cell (WBC) variants from (A). (D) Upset plot displaying the intersection of all plasma cfDNA variants from (A) across the different sequencing sources analyzed (plasma cfDNA, tumor tissue, matched WBC). Tumor tissue-specific variants were characterized from either whole exome sequencing of baseline tumor samples or clinical next-generation sequence data. All per-patient unique plasma variants are shown.

At first sampling, 24 of 26 patients had ≥1 tumor-derived ctDNA mutation, at a median mutant allele fraction (MAF) of 5.40% (range 0.11-19.70%, Supplementary Table S5). Evaluation of the landscape of tumor-derived mutations revealed frequent nonsynonymous alterations in TP53 (altered in 14 of 26 [54%] patients) and KRAS (9 of 26 [35%] patients). Tumor-derived mutations in ERBB2 (4/26 patients; 15%), ATM (2/26 patients; 8%), BRAF (2/26 patients; 8%), CDKN2A (2/26 patients; 8%) and CTNNB1 (2/26 patients; 8%) were also detected at >5% prevalence among patients (Fig. 2B). Comparison between the maximal MAFs of ctDNA mutations prior to therapy, revealed a difference in baseline MAFs between patients treated with immunotherapy and chemo-immunotherapy. For patients on single-agent immunotherapy, a lower cfTL at baseline was significantly associated with durable clinical benefit (Supplementary Fig. S2B, P = 0.029 Mann Whitney U test). However, patients on chemo-immunotherapy had similar distributions of ctDNA MAFs independent of therapeutic response (Supplementary Fig.S2B).

### ctDNA dynamics refine heterogeneous clinical responses

To evaluate changes in ctDNA dynamics under the selective pressure of therapy, we used the maximum MAF of tumor-derived sequence alterations detected at each timepoint as a measure of cfTL for each patient, and tracked cfTL during the course of single-agent immunotherapy or chemo-immunotherapy treatment. Using this approach, we were able to track cfTL dynamics for 26 patients with detectable ctDNA mutations in plasma and assign a molecular response classification. We defined molecular response (mR) as the complete clearance of cfTL following treatment initiation. Patients who displayed cfTL persistence across all sampled timepoints were assigned a classification of molecular progressive disease (mPD). A subset of cases displayed a transient molecular response followed by a subsequent increase in cfTL that was driven by either recrudescence of baseline ctDNA mutations or the emergence of mutations that were undetectable at baseline and were classified as molecular response followed by recrudescence or emergence.

Overall, 6 patients were classified in the molecular response category, with a time to molecular response as soon as 2 weeks from treatment initiation (median 4.5 weeks, range 2.29 – 14.86 weeks), while 13 patients were classified as molecular progressors (Supplementary Table S5, Fig. 3). Six patients had molecular response followed by recrudescence with a median time to molecular response of 7 weeks (range 3 – 33 weeks) (Supplementary Table S5, Fig.). One patient had molecular response followed by emergence (Supplementary Table S5, Fig. 3). Molecular response was not concordant with first radiographic response (P = 0.123, Fisher’s exact test), while a trend was noted towards a correlation between molecular response and best overall radiographic response (P= 0.056, Fisher’s exact test; Fig. 4A-C). To determine whether this limited concordance was attributed to the larger fraction of patients with radiographically stable disease in this cohort, we compared first and best overall radiographic response assessments and found that 3 patients with a first response assessment of stable disease were reclassified as partial responders at the time of best overall response assessment (Supplementary Table S1). cfTL analyses revealed that 2 of these 3 patients had a molecular response followed by recrudescence, with a time to molecular response as soon as 3 weeks (median 4.5 weeks, range 3 -6 weeks) (Supplementary Table S1). In these cases, ctDNA molecular response preceded best radiological assessments by an average lead time of 21 weeks (range 8 – 33 weeks). Conversely, 2 patients with a first radiographic response assessment of progressive disease were subsequently reclassified as stable disease at the time of best overall assessment. Analysis of ctDNA molecular response classifications revealed one patient to be a molecular progressor, consistent with short progression-free (PFS) and overall survival (OS) times of 1.4 months and 6.3 months, respectively. In contrast, the second patient was assigned a classification of molecular response followed by recrudescence, consistent with an extended PFS and OS of 23.0 months (Supplementary Table S1). Notably, molecular responses were concordant with durable clinical benefit (P=0.006, Fisher’s exact test, Fig. 4A, D), further supporting the accuracy of ctDNA molecular responses for capturing clinical outcomes. Collectively, these findings highlight the limitations of imaging-based response assessments for accurate evaluation of therapeutic benefit from immunotherapy and the association between ctDNA molecular response and durable therapeutic responses.

**Fig. 3.**
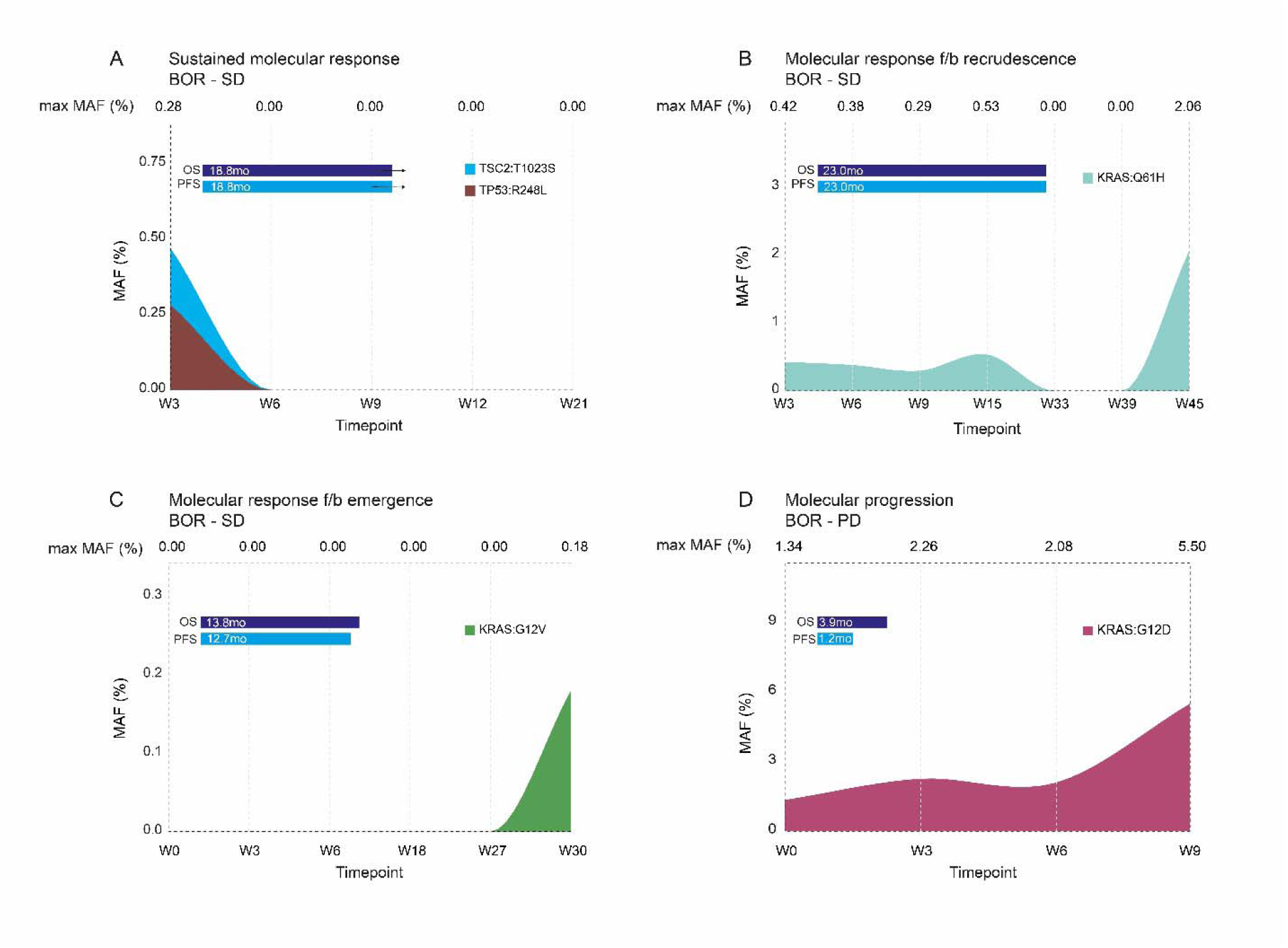
Longitudinal cfTL dynamics and molecular response classifications. (A) Dynamic changes in cfTL across serial plasma timepoints were analyzed for each patient and used to assign a molecular response classification. Representative examples of patients assigned to each response classification are shown. (A) Patient CGLU528 was classified as a molecular responder based on the complete clearance of cfTL from first sampling at week 3 to week 5 during first-line pembrolizumab treatment. This was consistent with the extended overall (OS) and progression-free (PFS) survival of this patient of 18.8 months. (B) Patient CGLU357 was assigned a classification of molecular response followed by recrudescence based on the complete clearance of cfTL from first sampling at week 3 to week 33, following completion of first-line pembrolizumab therapy. This was followed by a subsequent increase in cfTL at week 45 driven by the same KRAS Q61H variant detected at first sampling. The presence of cfTL clearance in this patient was consistent with an extended OS and PFS of 23 months. (C) Patient CGLU503 was assigned a classification of molecular response followed by emergence based on the absence of detectable cfTL from baseline to week 27 sampling during first-line carboplatin-pemetrexed-pembrolizumab treatment, which was followed by an increase in cfTL above the limit of detection at week 30, driven by the presence of a KRAS G12V variant. (D) Patient CGLU603 was classified as a molecular progressor based on the persistence of cfTL across all timepoints analyzed during first-line carboplatin-pemetrexed-pembrolizumab treatment. This was consistent with the short OS (3.9 months) and PFS (1.2 months) observed for this patient. BOR, best overall radiographic response; Max MAF, maximum mutant allele fraction.

**Fig. 4.**
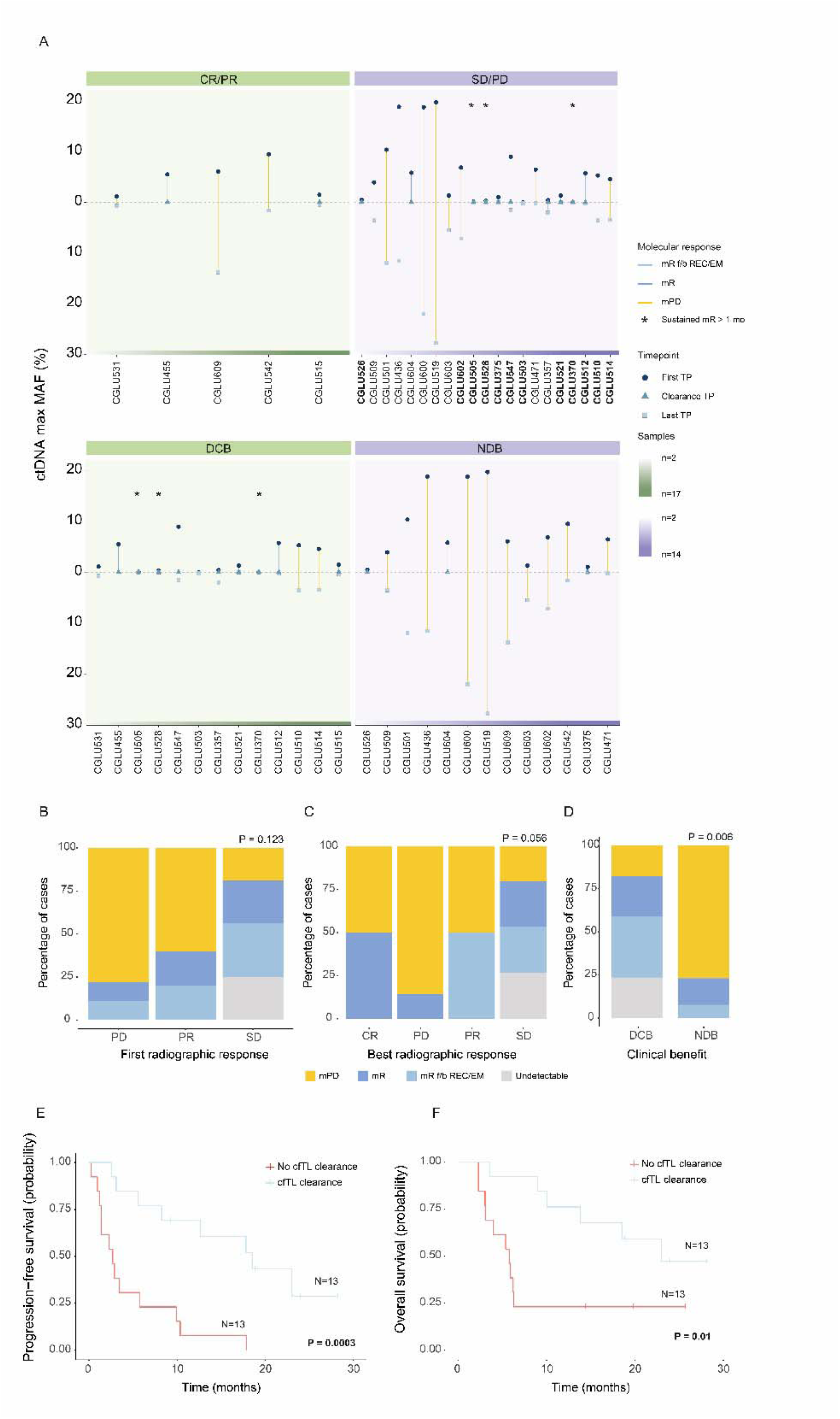
Association between ctDNA molecular responses and clinical outcomes. (A) Association between circulating tumor DNA (ctDNA) maximum mutant allele fraction (MAF) dynamics, first radiographic response classifications (top panels) and clinical benefit (bottom panels). Patients with a first radiographic assessment of stable disease (SD) are labelled in bold. The first plasma timepoint (TP) analyzed for each patient is shown as a navy circle and timepoints where cfTL clearance was first identified in plasma are shown as blue triangles. The final plasma timepoint analyzed for each patient is shown as a light-blue square if this was different from the timepoint of first clearance. Patients are ordered according to the total number of serial plasma samples available for analysis (indicated by gradient green and purple panels). Patients with a sustained molecular response > 1 month are indicated by an asterisk. (B-D) Whilst no associations were observed between ctDNA molecular responses and first radiographic response assessments (p = 0.123), molecular responses were trending in association with best overall radiographic responses (p=0.056) and were concordant with clinical benefit (p=0.006) for patients analyzed. Molecular response was more frequently observed among patients with durable clinical benefit compared to patients with non-durable clinical benefit. P values were calculated using Fisher’s exact tests. Patients with a molecular response defined by the presence of cfTL clearance across any timepoint analyzed had a significantly improved (E) progression-free (median PFS 18.51 vs 2.71 months) and (F) overall (median OS 23.01 vs 5.75 months) survival compared to patients with molecular progression.

Finally, we assessed whether ctDNA molecular responses were associated with overall (OS) and progression-free survival (Fig. 4E-F, Supplementary Fig.S3). Patients with molecular response or molecular response followed by either recrudescence or emergence had a significantly longer OS and PFS compared to individuals with molecular progression (median PFS 18.51 vs 17.82 vs 2.70 months, P = 0.001 and median OS 18.51 vs 23.01 vs 5.75 months, P = 0.04; Supplementary Fig.S3A-B). Stratification of patients according to the presence of molecular response across any timepoint analyzed confirmed these findings (median PFS of 18.51 vs 2.70 months, P = 0.0003 and median OS of 23.01 vs 5.75 months, P = 0.01 respectively; Fig. 4E-F). Furthermore, the relationship between molecular response and OS (HR = 0.15, 95% CI = 0.04-0.58, p = 0.006) and PFS (HR = 0.18, 95% CI = 0.06-0.56, p =0.003) was statistically significant after accounting for clinical covariates in a multivariate Cox proportional hazards regression model (Supplementary Tables S6-7).

Importantly, we repeated these analyses in the subset of 11 patients with a best overall radiographic response classification of stable disease and detectable cfTL and found that molecular response was significantly associated with PFS (median PFS 23.01 vs 5.75 months, P = 0.0075), with a trend towards longer OS (median OS 23.01 vs 6.28 months, P = 0.09) (Fig. 5). Collectively, these findings indicate the utility of cfTL monitoring and molecular response classifications for capturing the magnitude of therapeutic responses in patients with radiographically stable disease (Fig. 5).

**Fig. 5.**
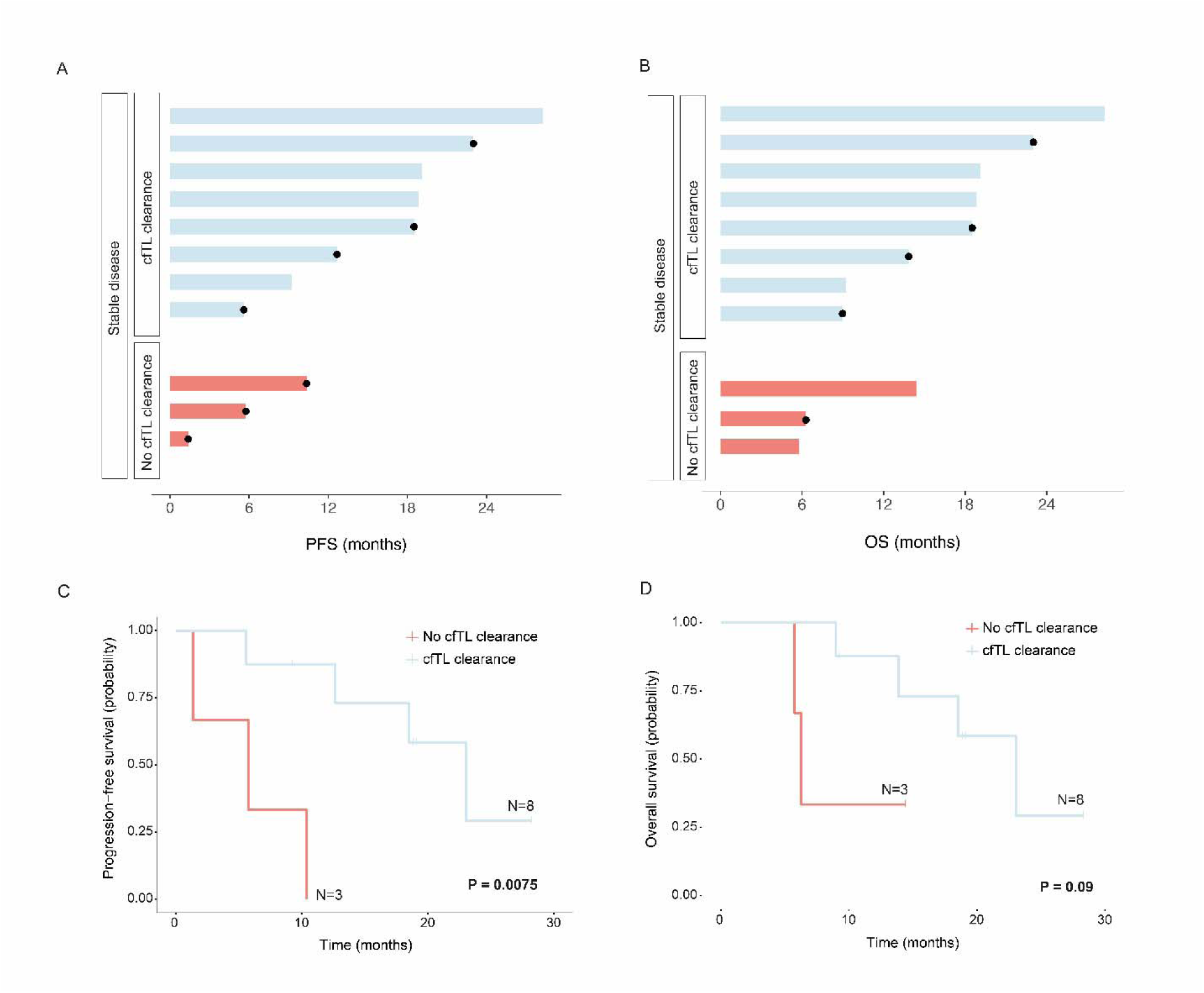
ctDNA molecular response is associated with survival outcomes in patients with radiographically stable disease. Swimmer plots showing (A) progression-free (PFS) and (B) overall (OS) survival of the subset of 11 patients in the cohort with a best overall radiographic response assessment of stable disease. Patients are stratified according to the presence (light blue) or absence (coral) of a ctDNA molecular response across plasma timepoints analyzed for the study. Death or disease progression are indicated by black circles. Molecular response, defined by the clearance of cfTL, was significantly associated with both (C) PFS (median PFS 23.01 vs 5.75 months) and (D) OS (median OS 23.01 vs 6.28 months) in these patients.

### Immune cell repertoire dynamics may point towards early emergence of immune-related toxicities

To assess the peripheral T cell repertoire in tumors responding to immune checkpoint blockade and evaluate differential dynamics associated with the development of irAEs, we performed TCR sequencing of serial buffy-coat samples collected at baseline (n=23) and during immunotherapy or chemo-immunotherapy treatment (n=58) from 30 patients. This included 23 patients who were also analyzed for cfTL dynamics in this study and 7 additional individuals from a previous cohort (Supplementary Table S8) (15). Paired analyses of peripheral TCR sequencing at baseline and on-treatment timepoints revealed no significant changes in clonotype abundances according to either molecular response or clinical benefit; however, significant on-treatment dynamic changes in circulating TCR clonotypes were observed in patients who experienced an irAE during treatment with checkpoint blockade compared to individuals who did not (Fig. 6A-E, Supplementary Figure S4, Supplementary Table S9). Clonotypes with a significant increase in abundance at on-therapy sampling compared to baseline were defined as expanding clones and those with a significant decrease in abundance following therapy initiation were defined as regressing clones (Methods). Patients who experienced an irAE harbored a higher fraction of both expanding and regressing TCR clones on-therapy, which was evident at the first timepoint sampled following treatment initiation (median 3 weeks, range 1 - 6.4 weeks) (p = 0.01 and p = 0.01, respectively; Fig. 6C-D, Supplementary Figure S4A-E), compared to individuals who did not develop an irAE. To characterize shared specificity amongst identified TCR clones, we performed clustering based on global similarity in CDR3 sequences, using the GLIPH2 method (Methods). Expansions of multiple TCR CDR3 clusters were observed in patients who experienced an irAE (Supplementary Tables S10-11). Of these, 15 clusters (g.GQGAYE, gD.GTE, gSLE.NSP, gSLGQ.SYE, gRAG.NTE, gSR.GSGANV, gS.TGQNTE, gS.TSGGDT, gSGG.NYG, gSL.GGSNQP, gSLLQG.E, gSLVGD, gSLVGG.YG, gSPGD.YE, gT.GGE) were enriched at baseline sampling in patients who experienced and irAE and were further clonally expanded on-therapy in these patients, compared to individuals who did not develop irAEs (Supplementary Table S11, Supplementary Fig.S5). Analyses of global cluster dynamics in individual patients revealed additional clusters that were significantly expanded on-therapy on a per-patients basis. As a representative example, TCR sequencing analysis from patient CGLU341 who experienced immunotherapy-related pneumonitis revealed both expanding and regressing TCR clones on-therapy (Fig. 6E), which were comprised of multiple global clusters. Significant increases in abundance were observed for gS.GQGYYG, gSS.GSP, gS.DRDYE, gS.QTNTE and gSLTGA.E clusters whilst gS.GLAGAYE, gS.RDNSP, gR.GVNTE and gSPG.G clusters were regressing on-therapy (Fig. 6F). Taken together, these findings suggest that TCR clonotypic dynamism in peripheral blood may be an indicator of emergence of irAEs during immunotherapy and highlight the potential for clustering of TCR CDR3 sequences to predict early onset of irAEs.

**Fig. 6.**
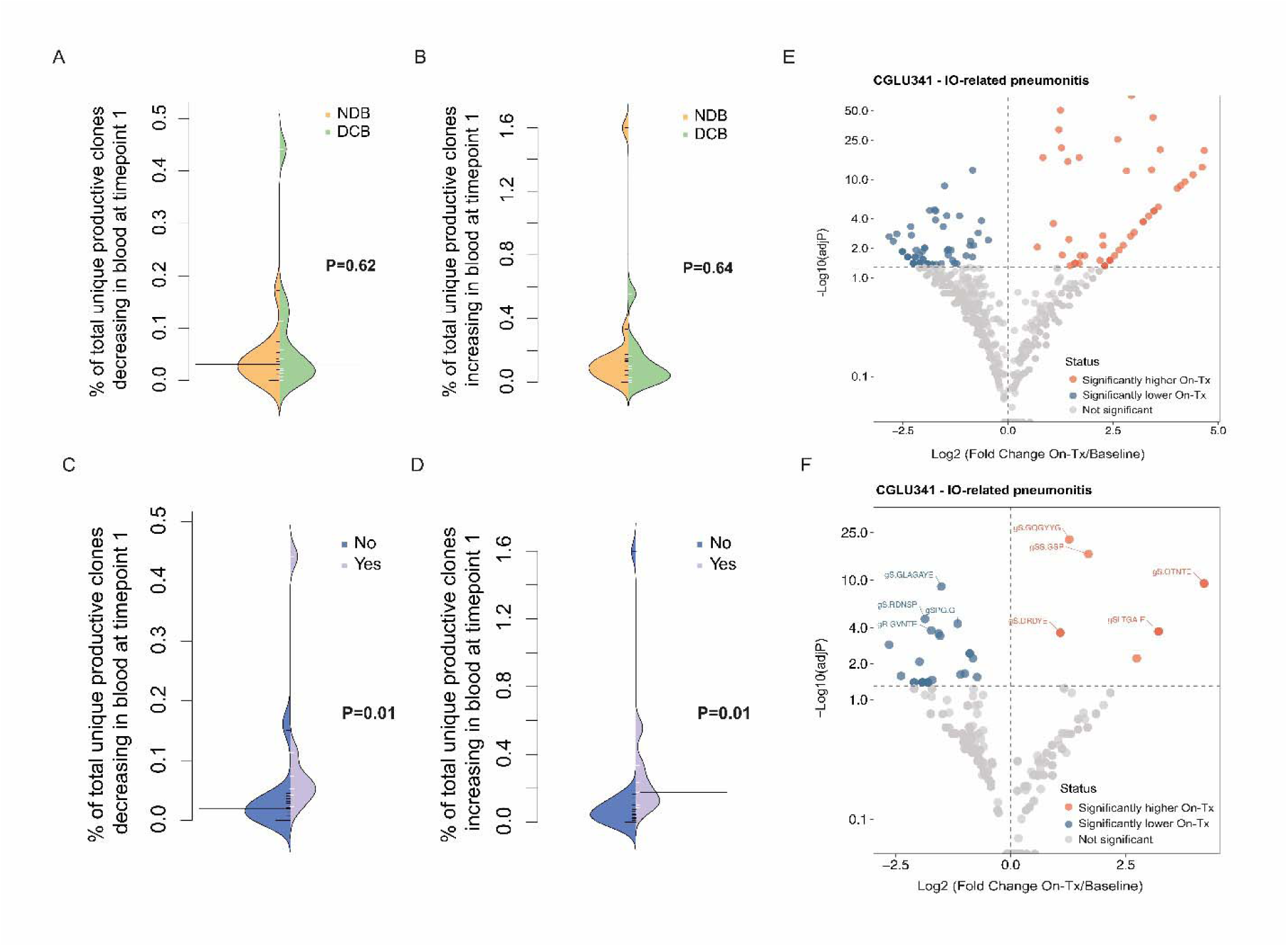
Peripheral TCR dynamics are associated with the development of immune-related adverse events (irAE) in patients treated with immunotherapy-containing regimens. (A, B) Whilst peripheral TCR clonotype dynamics were not associated with clinical benefit, (C, D) significant expansions and regressions of TCR clones were observed following the initiation of immunotherapy treatment in patients who developed an irAE (light purple), compared to individuals who did not (dark purple). The percentage of total unique productive TCR clones that were either (A, C) decreasing or (B, D) increasing in blood at the first timepoint analyzed during therapy compared to baseline are shown. (E) Volcano plot displaying TCR clonal dynamics in a patient (CGLU341) who developed immunotherapy-related pneumonitis. Clones that were significantly expanded at the first timepoint sampled on-therapy compared to baseline are shown in red and clones that regressed on-therapy are shown in blue. (F) TCR clones from (E) were clustered according to similarity in CDR3 sequences, using GLIPH2. TCR global clusters that were expanded on-therapy in patient CGLU341 are shown in red and clusters with a significantly lower abundance on-therapy compared to baseline are shown in blue. Global clusters with the greatest statistical significance according to the Fisher’s exact test are labelled.

To further investigate phenotypic differences in peripheral immune cell subsets that may be associated with the onset of irAEs, we evaluated serial plasma protein expression profiles from 28 patients. Comparison between baseline and on-therapy samples revealed increased expression of multiple mediators of adaptive immune responses on-therapy, including interferon (IFN)-gamma (P=0.0056, Mann Whitney U test), TNF (P=0.0003) and IL-10 (P=0.0001) cytokines, GZMA (P=0.001), GZMH (P=0.0003), the interferon stimulated chemokine CXCL9 (P < 0.0001) and PDCD1 (P < 0.0001) (Supplementary Table S12, Supplementary Fig.S6). Furthermore, in patients who experienced an irAE, we identified a significantly higher expression of pro-inflammatory mediators, including CCL19 (P=0.007), GZMA (P=0.017) and IL-6 (P=0.029), and T-cell markers, including CRTAM (P=0.017) and the T cell costimulatory molecule CD28 (P=0.017), at baseline sampling (Supplementary Table S12). Similarly, at on-therapy timepoints, patients who experienced an irAE had significantly higher expression levels of inflammatory markers, including IL-5 (P=0.04), CCL23 (P=0.01), MMP7 (P=0.005), ANGPT2 (P=0.02), and the inhibitory checkpoint PD-L2 (P=0.02), compared to individuals who did not develop toxicity (Supplementary Table S12). Collectively, our findings indicate that differences in the phenotypes of immune cell subsets, reflected in both baseline and on-therapy plasma protein expression profiles, may serve as sentinel features to identify individuals most likely to develop immunotherapy-related toxicities.

## Discussion

The challenges of predicting clinical response to immunotherapy in lung cancer mandate the development and clinical integration of minimally invasive real-time biomarkers to capture clinical response and guide clinical decision making (1-3,30,31). By analyzing cfTL dynamics during immunotherapy and chemo-immunotherapy in patients with advanced NSCLC, we identified that ctDNA molecular responses differentiated survival and could predict durable clinical benefit. The prevalence of cases with radiographically stable disease in this cohort provided a unique opportunity to assess the utility of molecular responses for characterizing the heterogenous nature of stable disease. Our results demonstrated that ctDNA molecular responses more accurately captured survival outcomes and the magnitude of therapeutic benefit in these cases, compared to imaging. Furthermore, our complementary assessment of the immune cell compartment revealed that peripheral blood T cell characteristics and plasma proteomic profiles are associated with the development of immune-related toxicities, with potentially important implications for clinical cancer care.

Our study provides support for the use of ctDNA molecular response as an early endpoint for therapeutic response to immunotherapy, using rigorous strategies to avoid non-tumor derived variants via matched WBC sequencing and informed by all available tumor tissue testing. While current commercial clinical liquid biopsy testing does not utilize either of these strategies consistently, incorporation of complementary sources of NGS sequencing has demonstrated benefit in NSCLC (32). Our study identified ctDNA molecular response, defined as the complete clearance of cfTL, as an important feature associated with survival and durable clinical benefit, highlighting the utility of ctDNA dynamics to guide clinical intervention for NSCLC. Importantly, our results indicate that ctDNA molecular responses may be particularly informative in patients with radiographically stable disease as the presence of a molecular response more accurately distinguished survival outcomes in these patients compared to imaging-based assessments.

Ongoing ctDNA-driven interventional clinical trials are investigating whether ctDNA molecular responses can be used to guide therapeutic change for patients with advanced or metastatic NSCLC (NCT04093167, NCT04166487, NCT04966676). The BR.36 clinical trial, in particular, specifically investigated the concordance between ctDNA molecular responses and radiographic responses (33). Similar applications of ctDNA-driven response assessments in the early-stage setting have shown potential for molecular responses in assessing the early on-therapy efficacy of neoadjuvant immunotherapy via tumor regression in patients with localized NSCLC (34). However, a unified definition of liquid biopsy-based molecular response is lacking in NSCLC and other solid cancers. On-treatment ctDNA decreases based on ctDNA clearance or ctDNA level reduction below given cutoffs have all been associated with clinical response (35–37). Molecular progression – or the increase or lack of significant decrease in ctDNA variants on-treatment – has potentially greater sensitivity in predicting clinical progression (8,15,36). Furthermore, the importance of time to molecular response assessment in the setting of immunotherapy – ranging from as early as three weeks (36) in NSCLC to up to twelve weeks (37) in various solid cancers – remains unsettled. Our findings suggest that on-treatment assessment for molecular response in both immunotherapy and chemo-immunotherapy, allow for timely adaptation to alternative therapies. However, our studies demonstrate that longitudinal on-treatment monitoring is necessary given the evidence of molecular recrudescence and emergence of new variants that can modify clinical outcomes.

In addition to the monitoring of clinical responses to immune checkpoint blockade, accurate and timely identification of immunotherapy-related toxicity to guide early intervention remains an unmet clinical need. Biological drivers underpinning the development of irAEs remain poorly characterized and there are no standard clinical biomarkers to identify high-risk patients. Beyond assessment of tumor response via ctDNA approaches, we demonstrated that TCR repertoire dynamics and plasma proteomic profiling are associated with immunotherapy-associated toxicity. Although these factors were not correlated with clinical benefit, we identified distinct associations between peripheral blood TCR clonotype dynamics and the development of irAEs. Clonal dynamics associated with irAE onset included the degree of T cell clonotypic expansions and regressions early during the course of treatment, which was driven by clusters of TCRs sharing similar CDR3 sequences and potential specificity for target antigens, consistent with the possibility of a self-reactive immunological mechanism underlying irAE development. Moreover, peripheral proteomic analysis revealed that elevated expression of proinflammatory cytokines and chemokine markers was further associated with irAE onset, consistent with a pattern of systemic immune activation. Further investigation of these results in larger pan-cancer cohorts are required to assess whether these features are able to distinguish between different irAE grades (18), determine the relative contributions of different immune cell subtypes to these dynamics and evaluate whether these findings generalize to the risk of immunotherapy-related toxicity in other solid tumor types.

Limitations of our study include the retrospective design and relatively small size of our study cohort. Furthermore, owing to the real-world nature of the study cohort, RECIST evaluations of radiographic response assessments were not available for patients on the study cohort.

Taken together, our study affirms the role of ctDNA-based molecular response as an independent predictor of clinical response with immunotherapy in NSCLC. Furthermore, we highlight the role of TCR sequencing and immunoproteomic profiling in assessing a patient’s risk of immunotherapy toxicity. Building on these findings, we anticipate that combinatorial liquid biopsy-based tests can aid clinical decision-making toward adaptive treatment regimens that improve patient outcomes, by rapidly and accurately capturing clinical outcomes while at the same time limiting untoward side effects.

## Acknowledgements

We would like to thank the Lung Cancer Precision Medicine Center of Excellence Investigators Sharon Penttinen, Michael Conroy, Durrant Barasa, Mary Anderson, Mariam Ghobadi-Krueger, Kenneth Harkness, Kerry Smith, Sreenivasa Idamakanti and Vivian Altiery De Jesus for their contributions. This work was supported in part by the US National Institutes of Health grants CA121113, CA062924 and CA006973 (V.A., R.B.S, and V.V.), the Bloomberg-Kimmel Institute for Cancer Immunotherapy (V.A., J.R.B and P.M.F.), the ECOG-ACRIN Thoracic Malignancies Integrated Translational Science Center grant UG1CA233259 (V.A., V.V.), the Emerson Collective Cancer Research Fund (V.A.), the V Foundation (V.A. and V.V.), the LUNGevity Foundation (V.A. and V.V.), the Hopkins-Allegheny Health Network (AHN) Cancer Research Fund (V.A. and A.Z.), the Conquer Cancer Foundation (J.C.M.), a MacMillan Scholars Award (J.C.M.), the Maryland Cigarette Restitution Fund (J.C.M.) and the International Lung Cancer Foundation (L.S.).

## Competing Interests Statement

J.C.M. has received research funding to Johns Hopkins University from the Conquer Cancer Foundation (Young Investigators Award) from Merck. V.A. receives research funding to Johns Hopkins University from Astra Zeneca and Labcorp/Personal Genome Diagnostics, has received research funding to Johns Hopkins University from Bristol-Myers Squibb and Delfi Diagnostics in the past 5 years and is an advisory board member for Neogenomics. V.A. is an inventor on patent applications (63/276,525, 17/779,936, 16/312,152, 16/341,862, 17/047,006 and 17/598,690) submitted by Johns Hopkins University related to cancer genomic analyses, ctDNA therapeutic response monitoring and immunogenomic features of response to immunotherapy that have been licensed to one or more entities. Under the terms of these license agreements, the University and inventors are entitled to fees and royalty distributions. R.B.S. is a founder of and holds equity in Delfi Diagnostics. He also serves as the head of data science for this organization. Johns Hopkins University owns equity in Delfi Diagnostics. This arrangement has been reviewed and approved by the Johns Hopkins University in accordance with its conflict-of-interest policies. S.C. has served in a consultant/advisory role for Genentech and Roche. J.F. has received research grants from Astra Zeneca, Pfizer and BMS and has served in a consultant role for Genentech, Eli lilly, AstraZeneca, Merck, Takeda, Coherus, Regeneron and Pfizer. K.M. has served in a consultant/advisory role for Amgen, Janssen, Mirati, AstraZeneca, Puma and has received research funding (directly to the institution) from Mirati and BMS. C.H. has served in a consultant/advisory role for AbbVie, Amgen, AstraZeneca, BMS, Genentech/Roche, Jannsen and GSK and has received research funding (directly to the institution) from AbbVie, Amgen, AstraZeneca, BMS, and GSK. B.L. has served in a consultant/advisory role for Janssen, Daiichi Sankyo, AstraZeneca, Eli Lilly, Genentech, Mirati, Amgen, Pfizer, BMS, Guardant 360 and Foundation Medicine. J.R.W is founder and owner of Resphera Biosciences LLC. P.F. has received research funding (directly to the institution) from AstraZeneca, BMS, Novartis, Regeneron, Kyowa and BioNTech, had served in a consultant/advisory role for AstraZeneca, Abbvie, Amgen, BMS, Novartis, Genentech, Sanofi, Surface, Janssen, G1 and Merck, and has served in a DSMB for Polaris and Flame. J.B. has received research funding from AstraZeneca and BMS, has served in a consultant/advisory role for Amgen, AstraZeneca, BMS, Genentech/Roche, Eli Lilly, GSK, Merck, Sanofi, Regeneron, Janssen and Johnson and Johnson, and has served in a DSMB for Janssen. V.L. has served in a consultant/advisory role for Takeda, Seattle Genetics, BMS, AstraZeneca and Guardant Health, and has received research funding from GSK, BMS, Merck and Seattle Genetics. J.N. has received research funding from Merck, AstraZeneca, BMS, Amgen, Genentech/Roche and Novartis, has served in a consultant/advisory role for Merck KGA, MSD, AstraZeneca, BMS, Genentech/Roche, Pfizer, Elevation Oncology, Takeda, Daiichi Sankyo, Kaleido Biosciences, NGM Pharmaceuticals, Amgen and Mirati, and has served in a DSMB for Daiichi Sankyo and AstraZeneca. V.E.V. is a founder of Delfi Diagnostics, serves on the Board of Directors and as a consultant for this organization, and owns Delfi Diagnostics stock, which is subject to certain restrictions under university policy. Additionally, Johns Hopkins University owns equity in Delfi Diagnostics. V.E.V. divested his equity in Personal Genome Diagnostics (PGDx) to LabCorp in February 2022. V.E.V. is an inventor on patent applications submitted by Johns Hopkins University related to cancer genomic analyses and cell-free DNA for cancer detection that have been licensed to one or more entities, including Delfi Diagnostics, LabCorp, Qiagen, Sysmex, Agios, Genzyme, Esoterix, Ventana and ManaT Bio. Under the terms of these license agreements, the University and inventors are entitled to fees and royalty distributions. V.E.V. is an advisor to Viron Therapeutics and Epitope. These arrangements have been reviewed and approved by the Johns Hopkins University in accordance with its conflict-of-interest policies. A.Z. serves on medical advisory boards and holds equity ownership of Previse, Gritstone Bio and Tg Therapeutics. A.L. is an employee of Delfi Diagnostics. J.P owns stock in Delfi Diagnostics. Under a license agreement between Delfi Diagnostics and the Johns Hopkins University, J.P. and the University are entitled to royalty distributions related to technology. This arrangement has been reviewed and approved by the Johns Hopkins University in accordance with its conflict-of-interest policies. K.M. receives research grants from Astra Zeneca, is a consultant for Pfizer, BMS, Roche, MSD, Abbvie, AstraZeneca, Diaceutics, Lilly, Bayer, Boehringer Ingelheim and Merck and has received honoraria from MSD, Roche, Astra Zeneca, Benecke. All other authors declare no conflicts of interest.

