## Supplemental Figures for "Elucidating the heterogeneity of immunotherapy response and immune-related toxicities by longitudinal ctDNA and immune cell compartment tracking in lung cancer"

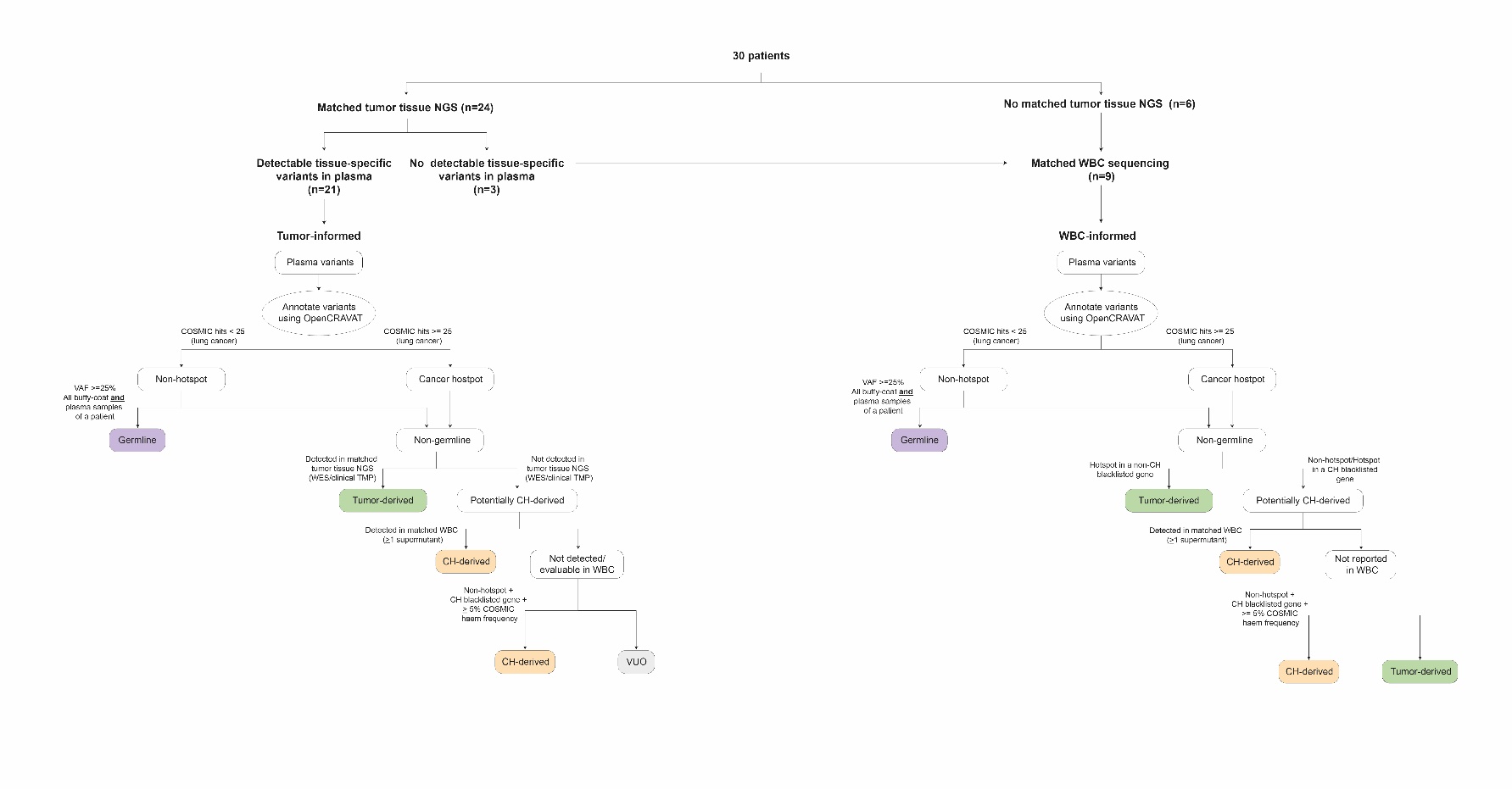
**Supplementary Fig.S1 Overview of branched logic architecture used for the classification of variant cellular origin in plasma.** Cell-free DNA (cfDNA) variants in plasma were assigned an origin classification of tumor-, germline- or clonal haematopoiesis (CH)-derived, using all available sequencing data. Patients with matched tumor tissue next-generation sequence (NGS) data available (n=24) from either whole exome sequencing or clinical tumor mutation profiling, were evaluated using our tumor-informed approach (Methods) for the identification of circulating tumor DNA (ctDNA) variants. As single tissue biopsies can be limited by sampling heterogeneity, patients with matched tumor NGS but no detectable tissue-specific variants in plasma (n=3) were further evaluated using our WBC-informed approach for identification of ctDNA variants missing by tissue sequencing. Patients with no detectable tumor-derived variants in plasma according to both approaches were subsequently classified as undetectable. Six patients without matched tumor tissue NGS available were evaluated using our WBC-informed approach for assignment of variant cellular origin (Methods). VUO, variant of unknown origin.

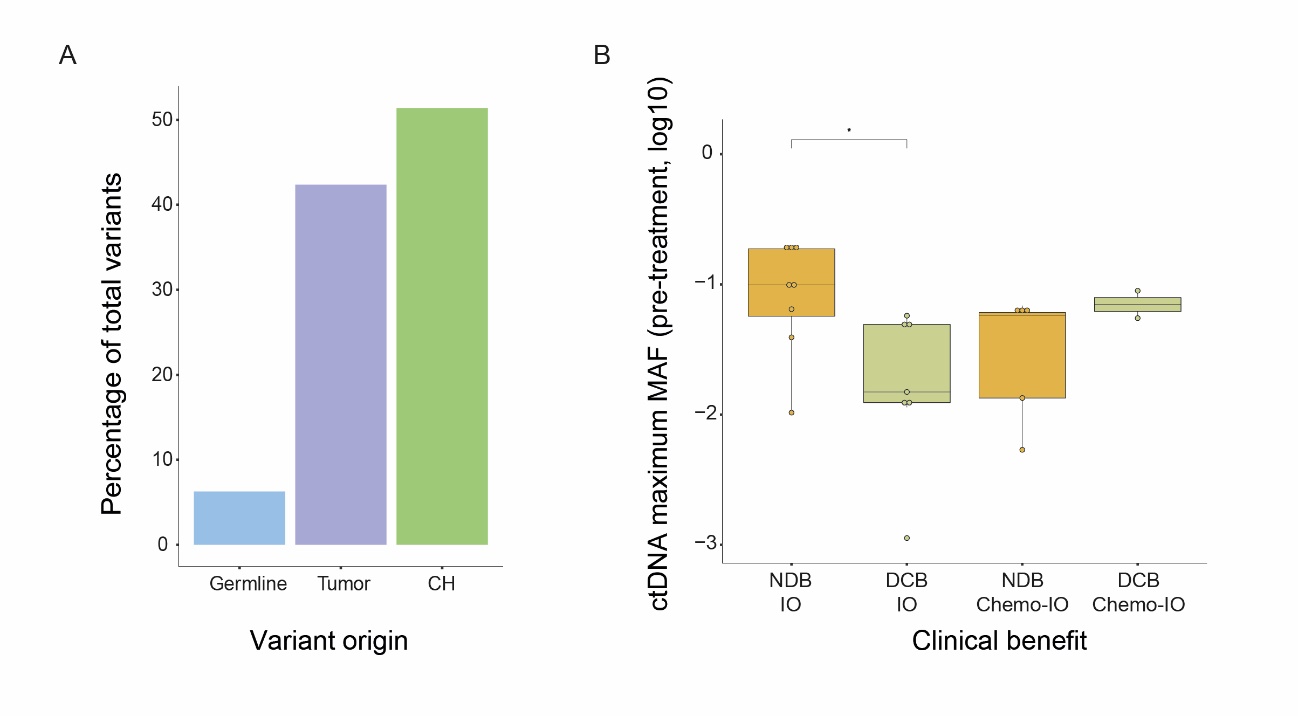

**Supplementary Fig.S2 Overview of variants detected in plasma cell-free DNA.** (A) The percentage of all classified plasma variants that were assigned an origin of germline, tumor-derived or clonal haematopoiesis (CH)-derived. (B) The maximum mutant allele fraction (MAF) of tumor-derived circulating tumor DNA (ctDNA) variants at baseline sampling, stratified by treatment (immunotherapy (IO) or chemo-immunotherapy (chemo-IO)) and clinical benefit (durable clinical benefit (DCB) or non-durable clinical benefit (NDB)). P values were calculated using the Mann-Whitney U test.

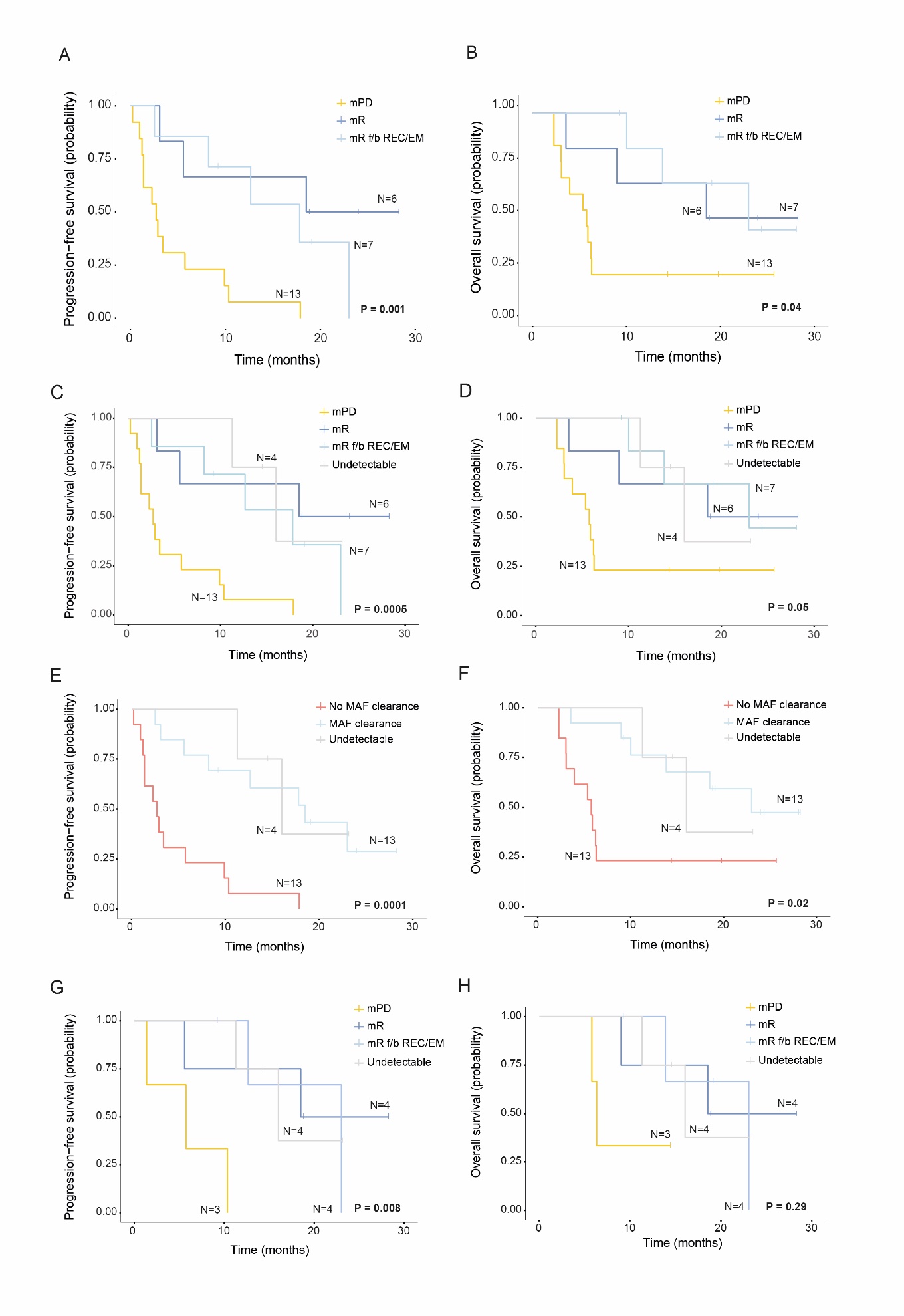

**Supplementary Fig.S3 Associations between ctDNA molecular responses and progression-free (PFS) and overall (OS) survival.** Molecular responses were significantly associated with both PFS (A) and OS (B) in all 26 patients with detectable cfTL (log-rank p value 0.001 and 0.04, respectively). Patients with a molecular response (mR) are shown in *blue*, patients with a molecular response followed by either recrudescence or emergence (mR f/b REC/EM) are shown in *light blue* and patients with molecular progression (mPD) are shown in *yellow*. (C, D) Analyses shown in (A) and (B) were repeated with the subset of patients (n=4) who were classified as having undetectable cfTL (*grey*). (E, F) Associations between the presence *(light blue)* or absence *(coral)* of a molecular response, defined by the presence of cfTL clearance across any sampled timepoint, and survival outcomes, including cases with undetectable cfTL *(grey)*. (G, F) Associations between molecular responses and survival outcomes in patients with a best overall radiographic response assessment of stable disease.

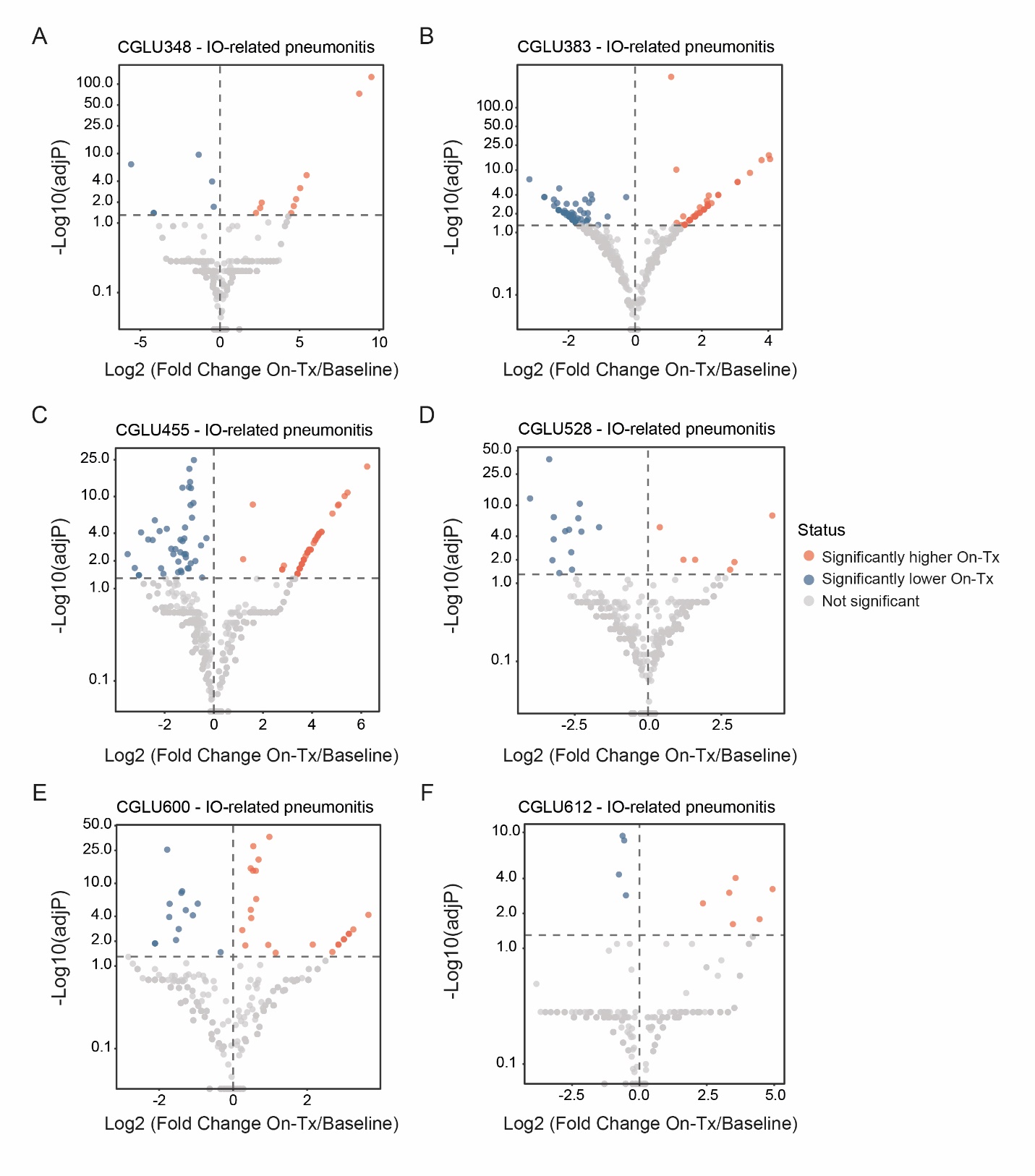

**Supplementary Fig.S4 T cell receptor (TCR) clonotype dynamics in patients who developed immunotherapy-related toxicity.** (A-F) Volcano plots showing significant TCR clonotypic expansions *(red)* and regressions *(blue)* following treatment with immunotherapy-containing agents in patients who developed immunotherapy (IO) – associated pneumonitis. On-Tx, On-treatment.

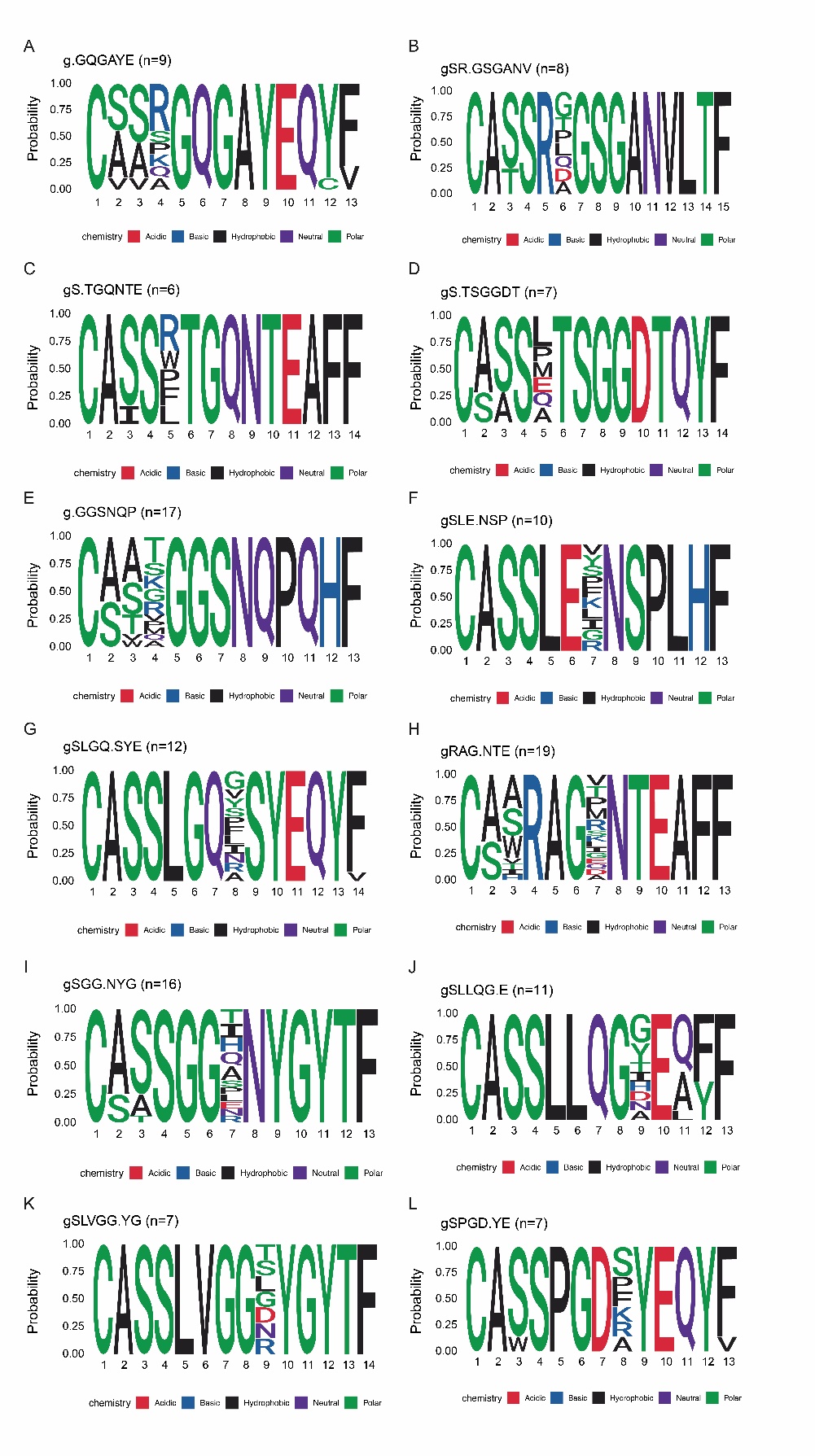
**Supplementary Fig.S5 Global TCR clusters that were enriched in patients who developed immunotherapy-related toxicity.** (A-L) Logo plots showing examples of global TCR clusters identified using GLIPH2 that were enriched at baseline were further significantly expanded on-therapy in patients who developed an immune-related adverse event (irAE), compared to individuals who did not. The size of each cluster, defined by the number of distinct CDR3 member sequences, is indicated alongside each panel. Enriched clusters with a minimum motif length of 6 are shown.

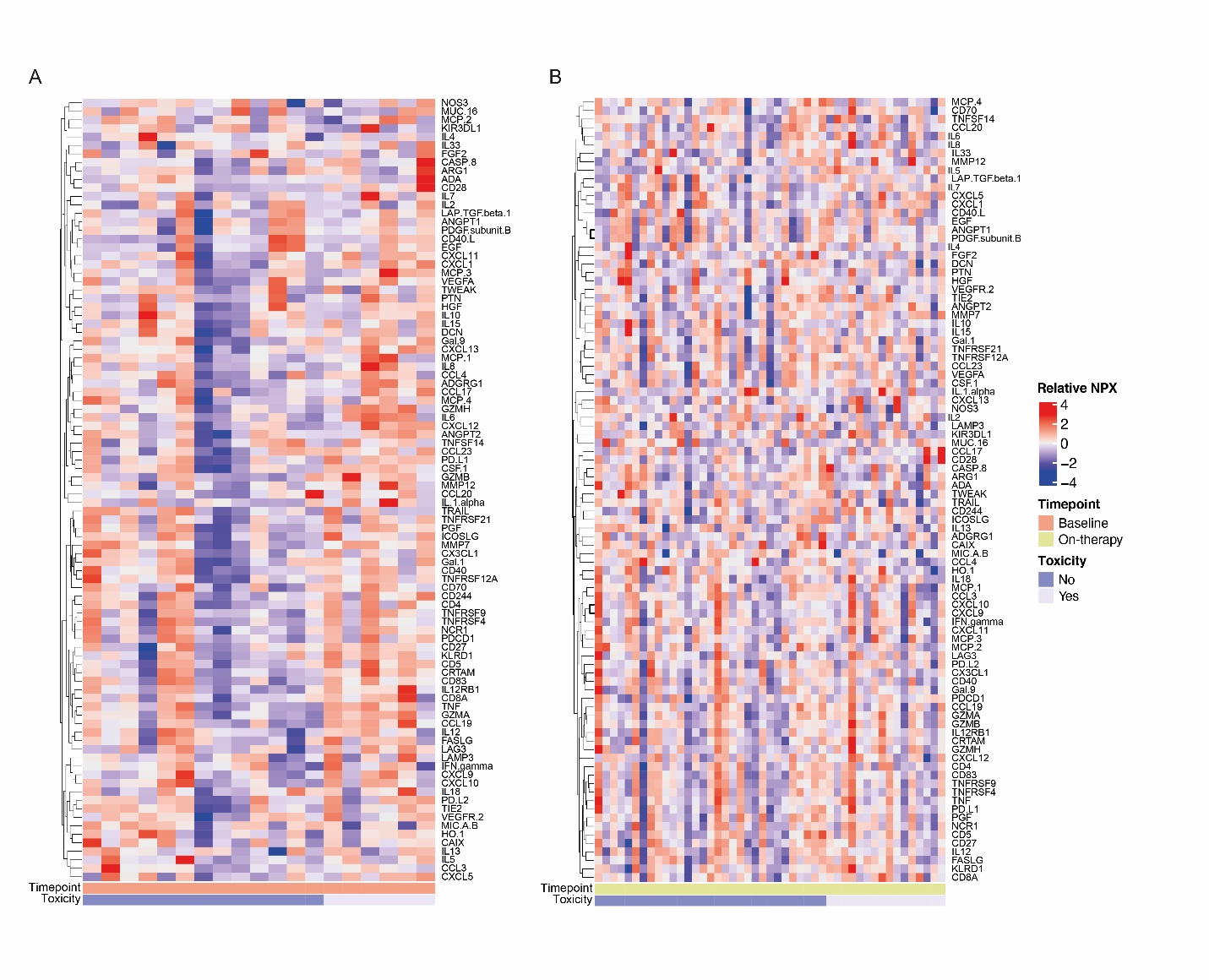

**Supplementary Fig.S6 Comparison between plasma proteomic profiles across treatment timepoints and toxicity.** Heatmaps showing normalized protein expression (NPX) scores for 92 proteins evaluated using Olink analyses across all (A) baseline (n=19) and (B) on-therapy (n=47) plasma samples from 28 patients. Panels indicating treatment timepoint and the presence of absence of immunotherapy-related toxicity are displayed below the heatmaps.
